# Alternative splicing of GluN1 gates glycine-primed internalization of NMDA receptors

**DOI:** 10.1101/2020.12.18.423454

**Authors:** Hongbin Li, Vishaal Rajani, Lu Han, Danielle Chung, James E. Cooke, Ameet S. Sengar, Michael W. Salter

## Abstract

N-methyl-D-aspartate receptors (NMDARs), a principal subtype of excitatory neurotransmitter receptor, are composed as tetrameric assemblies of two glycine-binding GluN1 subunits and two glutamate-binding GluN2 subunits. Gating of the NMDARs requires binding of four co-agonist molecules, but the receptors can signal non-ionotropically through binding of glycine, alone, to its cognate site on GluN1. A consequence of this signalling by glycine is that NMDARs are primed such that subsequent gating, produced by glycine and glutamate, drives receptor internalization. The GluN1 subunit is not a singular molecular species in the CNS, rather there are 8 alternatively spliced isoforms of this subunit produced by including or excluding the N1 and the C1, C2 or C2’ polypeptide cassettes. Whether alternative splicing affects glycine priming signalling is unknown. Here, using recombinant NMDARs expressed heterologously we discovered that glycine priming of NMDARs critically depends on alternative splicing: the four splice isoforms lacking the N1 cassette, encoded in exon 5, are primed by glycine whereas glycine priming is blocked in the four splice variants containing the N1 cassette. On the other hand, the C-terminal cassettes – C1, C2 or C2’ – had no effect on glycine priming signalling. Nor was glycine priming affected by the GluN2 subunit in the receptor. In wild-type mice we found that glycine primed internalization of synaptic NMDARs in CA1 hippocampal pyramidal neurons. With mice we engineered such that GluN1 obligatorily contained the N1 cassette, glycine did not prime synaptic NMDARs in pyramidal neurons. In contrast to pyramidal neurons, we discovered that in wild-type mice, synaptic NMDARs in CA1 inhibitory interneurons were resistant to glycine priming. But we recapitulated glycine priming in inhibitory interneurons in mice engineered such that GluN1 obligatorily lacked the N1 cassette. Our findings reveal a previously unsuspected molecular function for alternative splicing of GluN1 in controlling non-ionotropic signalling of NMDAR by glycine and the consequential cell surface dynamics of the receptors.

## INTRODUCTION

*N*-Methyl-D-aspartate receptors (NMDARs) are heterotetrameric ionotropic receptors found throughout the central nervous system (CNS), and play critical roles in neuronal development, synaptic plasticity and disease[1, 2]. NMDARs are assembled as two heterodimers each composed of a glycine-binding GluN1 subunit and a glutamate-binding GluN2 subunit. GluN1 is encoded by a single gene, with 8 splice variants, whereas there are four GluN2 genes, encoding subunits GluN2A-D [3]. Each of the GluN1 and GluN2 variants are competent to generate functional heterotetramers, yielding considerable diversity to the composition of NMDARs across the CNS.

Transcription and processing of *GRIN1*, the gene encoding GluN1, produces the eight mRNA variants through inclusion or exclusion of exons 5 and 21 and incorporating either exon 22 or exon 22’ (Fig. 1A). The polypeptide cassettes encoded by these exons are referred to as N1, C1, C2 and C2’, respectively [4, 5]. The GluN1 splice isoforms are referred to as follows [3]: a, N1-lacking; b, N1-containing; 1-1, C1- and C2-containing; 1-2, C1-lacking/C2-containing; 1-3, C1- and C2’-containing; 1-4, C1-lacking/C2’-containing (Fig. 1A). GluN1 splice variants are differentially expressed in various types of neurons, across different brain regions, and throughout distinct development stages [6–9]. N-terminal splicing affects NMDAR channel gating kinetics[10–12], as well as modulation by pH, Zn^2+^, extracellular polyamines [13] and long-term synaptic potentiation [14]. Many important properties of NMDA receptors such as glycine and glutamate potency, and voltage-dependent blockade by Mg^2+^ are unaffected by the presence of N1 insert[4, 5, 15]. C-terminal variants are implicated in processing NMDARs in the endoplasmic reticulum, Golgi and export to the cell surface [16–20].

**Figure 1.**
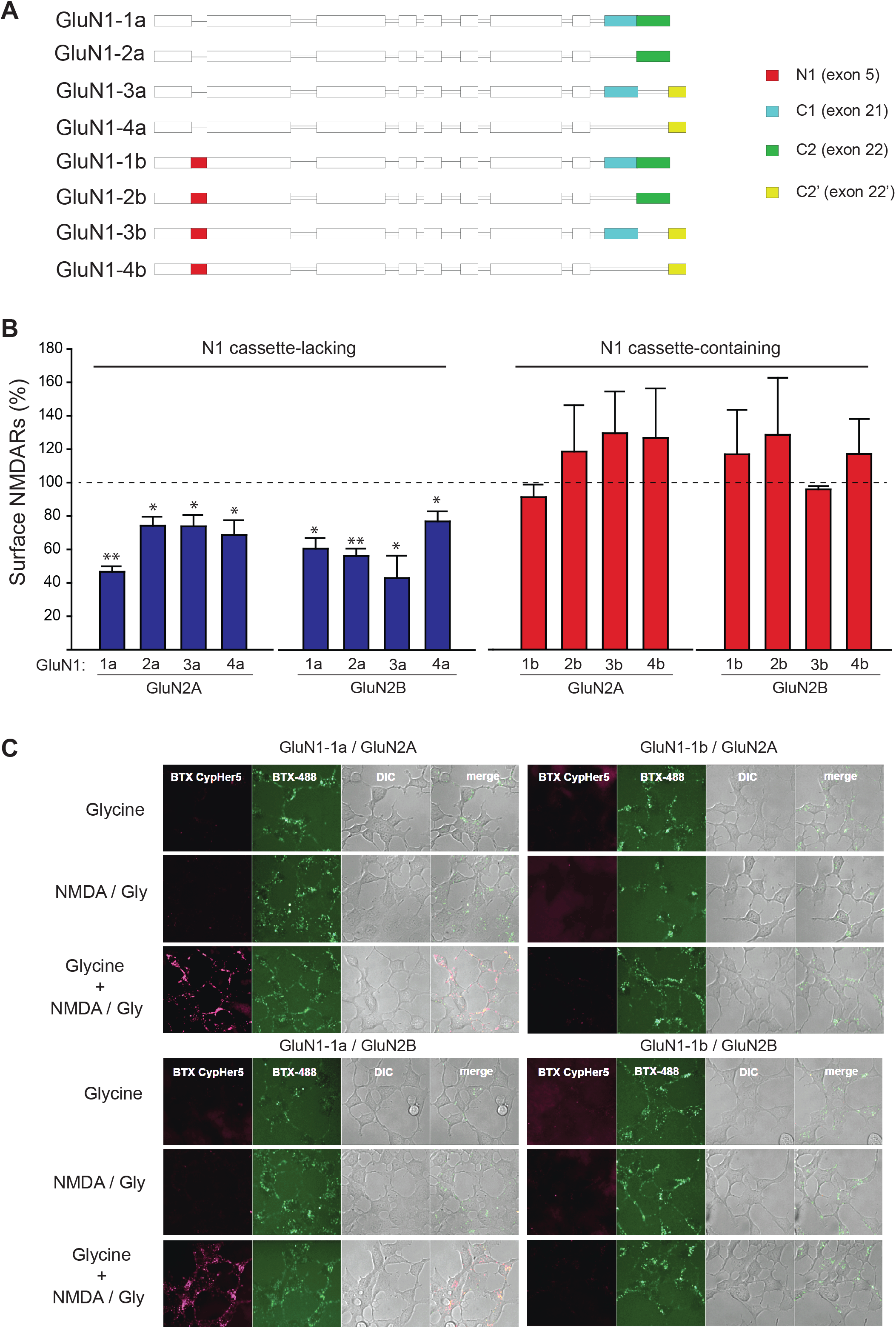
GluN1 splice variants regulate glycine-primed NMDAR internalization in HEK-293 cells. **(A)** Schematic of the 8 alternative splice variants of GluN1. **(B)** Cell surface expression of NMDARs expressing each GluN1 splice variant with either GluN2A or GluN2B quantified by cell ELISA assay from HEK293 cells after receptor priming (Gly 100 μM, 5 min) and activation (NMDA 50 μM, Gly 1 μM, 5 min). Data are grouped by inclusion of the N1 cassette, and then by GluN2 subunit. Data are normalized to glycine primed levels prior to receptor activation. The surface NMDARs composed of GluN1 subunits lacking the N1 cassette decreased to a range from 42.9 ± 13.1% to 76.8 ± 5.2% of unprimed NMDARs whereas the surface NMDARs composed of GluN1 subunits containing N1 cassette remained within the range of 91.2 ± 6.8 % to 129.5 ± 24.3% of unprimed NMDARs. Statistical significance is indicated with *p < 0.05, **p < 0.01 by Student t-test. **(C)** Representative confocal microscopy images of HEK293 cells expressing recombinant NMDARs with either BBS-tagged GluN1-1a or GluN1-1b, expressed with either GluN2A or GluN2B. Internalized NMDAR receptors are visible in the red channel (BTX-Cypher5E) versus total NMDAR expression in the green channel (BTX-AF488). Differential interference contrast (DIC) images reveal the number of cells in the corresponding field of view.

Canonical signalling by NMDARs is mediated by its ionotropic function initiated through simultaneous binding of two molecules of each of the co-agonists glycine (or D-serine) and glutamate to the ligand-binding domains in extracellular regions of the receptor which produces conformational changes that open the cationic conductance pathway of the receptor complex[21, 22]. However, increasing evidence is accumulating for non-canonical signalling by NMDARs initiated by independent binding of glycine or glutamate, which does not cause opening of the ionic conductance pathway but which nevertheless causes conformational changes that are transmitted across the membrane resulting in molecular rearrangements within the cell [23]. A striking example of non-ionotropic signalling by NMDARs was the discovery [24] that glycine binding to its cognate extracellular site, without binding of glutamate to its cognate site, drives intracellular recruitment of the AP2 endocytic adaptor complex. AP2 association primes NMDARs for subsequent internalization initiated by simultaneous binding of both co-agonists glycine and glutamate.

The molecular constraints within NMDARs for the non-canonical signalling by glycine remain unknown. Glycine-primed enhancement of AP2 association and subsequent internalization has been demonstrated with recombinant NMDARs expressed heterologously [24] which has led to the identification of one amino acid residue in GluN1 critical for priming by glycine [25]. This residue is 100% conserved across species. Given the natural diversity of GluN1 isoforms in the CNS and linkage of splicing to disease [26] a key open question remains as to whether alternative splicing of *GRIN1* affects glycine-priming of NMDARs. Therefore, here we investigated whether GluN1 splice variants may play a crucial role in the GluN1 signal transduction of glycine-primed internalization. We examined glycine priming with all 8 splice isoforms with heterologously expressed NMDARs, and then with representative isoforms in mice engineered to express only GluN1a and GluN1b variants.

## RESULTS

### N1 cassette in GluN1 prevents glycine-primed internalization of recombinant NMDARs

To investigate whether alternative splicing of *Grin1* affects glycine primed internalization of NMDARs we co-transfected HEK293 cells with one of the eight GluN1 splice variants together with either GluN2A or GluN2B. We verified that each of the constructs led to expression of protein which was trafficked to the cell surface (Supplementary Fig. 1). NMDARs were probed for glycine-primed internalization in two steps. First, the cells were conditioned with extracellularly applied glycine (100 μM), a concentration we previously found to prime subsequent internalization of GluN1-1a containing NMDARs [24, 25]. Second, the cells were treated with control extracellular solution (ECS) or with ECS containing NMDA (50 μM) plus glycine (1 μM) (NMDA+glycine treatment). Cells were fixed without permeabilization and the cell surface expression of NMDARs was quantified by enzyme-linked immunosorbent assay (ELISA). For NMDARs expressing GluN1 isoforms lacking the N1 cassette (i.e. the ‘a’ isoforms) we found that after conditioning glycine and NMDA+glycine treatment the surface level was significantly less than the surface level after conditioning glycine and treating with ECS (Fig. 1B). The glycine-primed decrease in NMDAR surface level of the ‘a’ isoforms was observed irrespective of intracellular GluN1 C-tail variants or whether GluN1 was co-expressed with GluN2A or with GluN2B (Fig. 1B). By contrast, the surface levels of NMDARs expressing the ‘b’ isoforms were not affected by conditioning glycine and NMDA+glycine treatment regardless of the GluN1 C-tail variants or co-expression with GluN2A or GluN2B. Thus, there was a striking difference between NMDAR isoforms lacking the N1 cassette, all of which showed glycine-primed internalization, as compared with isoforms containing the N1 cassette in GluN1, none of which showed glycine-primed internalization. Given this difference, we focused our subsequent investigations with recombinant NMDARs on those composed of GluN1-1a or GluN1-1b as representative of receptors, lacking or containing, respectively, the N1 cassette of the GluN1 subunit.

As an approach to test for glycine-primed internalization in addition to ELISA, we visualized changes in NMDAR surface expression by expressing GluN1 subunits in which we had inserted a bungarotoxin binding site (BBS) into the N-terminus of each isoform. We have previously established for NMDARs containing GluN1-1a that BBS-labelling has no effect on cell surface expression or function [25]. At the beginning of each experiment, we labelled cell surface BBS-NMDARs with α-bungarotoxin (BTX) conjugated with CypHer5E, a dye which is fluorescent only in acidic pH [27], as found in the lumens of endosomes. Then at the end of the experiment, we labelled BBS-NMDARs with BTX conjugated Alexa Fluor 488 (BTX-AF488) to visualize NMDARs remaining the cell surface. In cells only conditioned with glycine or only treated with NMDA+glycine we found no CypHer5E fluorescence above background whereas AF488 was readily detected on cells regardless of NMDAR composition (Fig. 1C). Thus, neither glycine alone nor NMDA+glycine alone caused endocytosis for GluN1-1a or GluN1-1b with either GluN2A or GluN2B subunits. On the other hand, bright CypHer5E fluorescence was observed in cells expressing GluN1-1a, together with GluN2A or GluN2B, following conditioning with glycine and treatment with NMDA+glycine, indicating that NMDAR endocytosis had occurred (Fig. 1C, left lower panels). However, in cells expressing GluN1-1b, conditioning with glycine and treatment with NMDA+glycine failed to drive Cypher5E fluorescence, with either GluN2 subunit, (Fig. 1C). Thus, our observations from imaging were consistent with those from ELISA, and from these convergent findings together we conclude that the presence of N1 cassette in GluN1 subunits prevents glycine-primed NMDAR internalization, i.e. the absence of the N1 cassette permits NMDAR internalization upon conditioning with glycine and treatment with NMDA+glycine.

### N1 cassette prevents glycine-primed depression of recombinant NMDAR currents

We previously observed with recombinant NMDARs containing GluN1-1a that conditioning glycine causes a progressive, use-dependent decline of NMDA-evoked currents [24, 25]. To examine the effect of the N1 cassette of GluN1 on NMDAR currents following glycine conditioning, we performed whole-cell patch-clamp recordings from HEK293 cells co-transfected with GluN1-1a or GluN1-1b together with either GluN2A or GluN2B subunits. NMDAR currents were evoked by applying NMDA (50 μM) plus glycine (1 μM) at 60 second intervals (Fig. 2). After a stable baseline current response had been established, the NMDA+glycine applications were temporarily suspended during at which time glycine in ECS or ECS was applied. The competitive glutamate-site antagonist D-APV (100 μM) was included to prevent the possibility of NMDA channel gating. We found that for the first several applications after resuming the NMDA+glycine applications, the amplitude of NMDAR currents in cells conditioned with glycine was not different from that in cells receiving ECS. Subsequently, in cells expressing NMDARs containing GluN1-1a with either GluN2A or GluN2B and conditioned with glycine+D-APV we found that NMDAR currents progressively decreased in amplitude. The level of the currents 20 minutes after conditioning was significantly less than that of NMDAR currents from cells conditioned with ECS+D-APV without glycine (Fig. 2A,B). By contrast, in cells expressing GluN1-1b together with GluN2A or with GluN2B, applying glycine+D-APV did not cause a significant decline in evoked current versus applying the ECS+D-APV control (Fig. 2C,D). Taking these findings together, we conclude that inclusion of the N1 cassette in GluN1 prevents the glycine-primed decrease of NMDAR-mediated currents.

**Figure 2.**
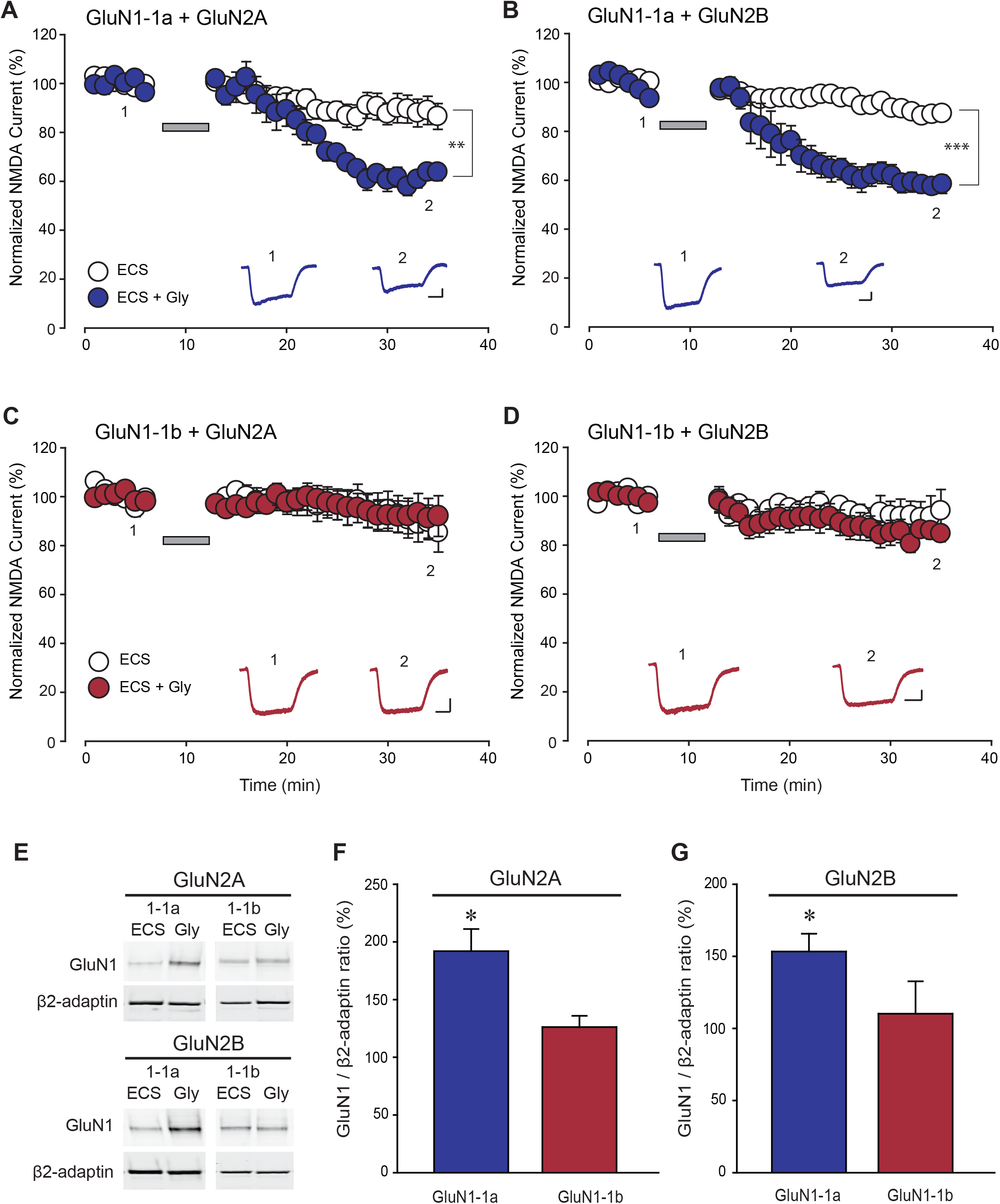
N1 cassette prevents the glycine priming-induced decrease of NMDAR current in HEK293 cells. Time series of normalized NMDAR peak current evoked by 50 μM NMDA and 1μM glycine following treatment with ECS containing 100 μM glycine and D-APV (filled circles) or ECS containing D-APV (open circles) from HEK293 cells expressing **(A)** GluN1-1a/GluN2A receptors (58.9 ± 3.8% vs 88.4 ± 5.8 %, n = 6 & 7 respectively, **p < 0.01); **(B)** GluN1-1a/GluN2B receptors (58.1 ± 3.8% vs 87.3 ± 2.5%, n = 5 & 7 respectively, ***p ≤ 0.001); **(C)** GluN1-1b/GluN2A receptors (91.3 ± 8.6%, vs 88.6 ± 8.0, n = 4 & 5 per group) and **(D)** GluN1-1b/GluN2B receptors (84.8 ± 3.5% vs 92.5 ± 8.6%, n = 5 & 4 respectively). Bath application is denoted by the grey bar. At the bottom of each panel are representative averaged traces from five consecutive evoked currents at the indicated times (1 and 2). **(E)** Representative immunoblot of GluN1 and β2-adaptin with immunoprecipitation of β2-adaptin from HEK293 cells expressing GluN1-1a or GluN1-1b together with GluN2A (top) or GluN2B (bottom) after conditioning with ECS + D-APV or ECS + glycine + D-APV. **(F-G)** Histogram of averaged GluN1/β2-adaptin ratio following conditioning with 100 μM glycine and D-APV in HEK293 cells expressing (F) GluN1-1a/GluN2A (191.9 ± 19.2%) vs GluN1-1b/GluN2A (126.1 ± 9.8%, *p < 0.05) and (G) GluN1-1a/GluN2B (153.3 ± 12.3%) vs GluN1-1b/GluN2B (110.2 ± 22.5%, *p < 0.05). Student t-test is used for all statistic comparisons.

### N1 cassette prevents glycine-primed recruitment of AP2 to recombinant NMDARs

A hallmark of glycine priming of NMDARs is recruitment of the AP2 protein complex upon glycine stimulation which readies the receptor for subsequent internalization[24]. Therefore, we investigated whether including the N1 cassette of GluN1 affects glycine-stimulated recruitment of AP2. To this end we immunoprecipitated the β2 subunit of AP2 from HEK293 cells expressing GluN1-1a or GluN1-1b together with GluN2A or GluN2B after conditioning with glycine+D-APV or only D-APV in ECS (Fig. 2E-G). With cells expressing GluN1-1a, conditioning with glycine led to a significant increase in the amount of this GluN1 subunit co-immunoprecipitating with AP2. Conditioning with glycine increased co-immunoprecipitation of GluN1-1a in cells co-transfected with GluN2A or with GluN2B. However, glycine conditioning did not change the amount of GluN1-1b-containing NMDARs co-immunoprecipitating with AP2 (Fig. 2E-G).

Overall, for recombinantly expressed NMDARs, we found that including the N1 cassette in GluN1 prevents: i) glycine-primed loss of NMDARs from the cell surface, ii) glycine-primed internalization of NMDARs into acidic organelles, iii) glycine-primed decline of NMDAR currents, and iv) glycine-stimulated recruitment of AP2 to the NMDAR complex. That is, the indicia of glycine priming [28] are prevented by the N1 cassette of GluN1 in heterologously expressed NMDARs activated by exogenous agonists.

### Glycine primes a decline in synaptic NMDAR currents in hippocampal CA1 pyramidal neurons from rats and mice

To assess the effect of glycine on native NMDARs activated endogenously, we investigated NMDAR excitatory postsynaptic currents (EPSCs) in CA1 pyramidal neurons in *ex vivo* hippocampal slices. NMDAR EPSCs were pharmacologically isolated by blocking AMPA and GABAA receptors with bath applied CNQX and bicuculline, respectively. Synaptic responses were evoked by stimulating Schaffer collateral inputs every 10 seconds; NMDAR EPSCs were recorded with the membrane potential held at +60 mV. After establishing a stable baseline amplitude of NMDAR EPSCs, we conditioned the slices by bath applying glycine in artificial cerebrospinal fluid (ACSF) or ACSF alone (e.g. Fig. 3A). In recordings from pyramidal neurons in slices from Sprague Dawley rats, we found that just after resuming the Schaffer collateral stimulation the amplitude of NMDAR EPSCs in cells conditioned with glycine had increased slightly, as compared with the baseline level, but was indistinguishable from the amplitude of NMDAR EPSCs in cells conditioned with ACSF (Fig. 3A). However, during the recordings, glycine conditioning subsequently led to a gradual decrease in NMDAR EPSC amplitude to a level significantly less than that observed with conditioning with ACSF alone (Fig. 3A,B). To test whether the decline in NMDAR EPSCs was mediated by dynamin [28] we made recordings in which the intracellular solution was supplemented with a dynasore (50 μM), a small molecule inhibitor of dynamin [29]. We found that during recordings in which dynasore was included in the intracellular solution, conditioning with bath-applied glycine did not induce the decline in NMDAR EPSC amplitude observed in recordings lacking intracellular dynasore (Fig 3A,B).

**Figure 3.**
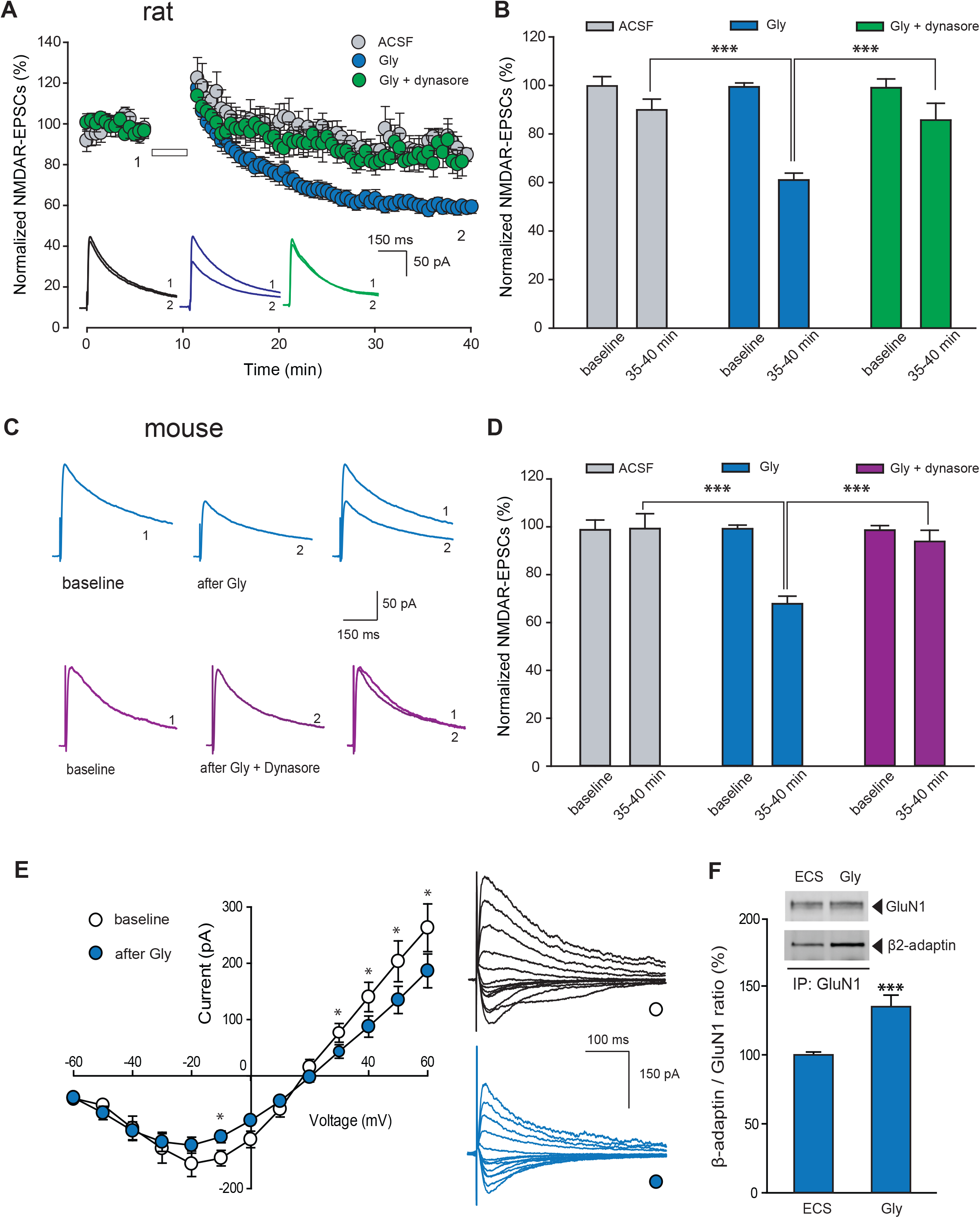
Glycine-primed decline in synaptic NMDAR current is dynamin dependent in rats and mice. **(A)** Scatter plot of NMDAR EPSC peak amplitude over time from rat CA1 pyramidal neurons recorded following 5 min of bath-applied ACSF (grey), glycine (0. 2 mM, blue) or glycine (0.2 mM) with intracellularly applied dynasore (50 μM, green). Bath application is denoted by the white bar. Representative average NMDAR EPSC traces were recorded at membrane potential of +60 mV at the times indicated (1 and 2). **(B)** Histogram of averaged NMDAR EPSCs measured before (baseline) and after treatment of ACSF (89.9 ± 4.4%, n = 6, grey), glycine (61.0 ± 2.9%, n = 12, blue, ***p ≤ 0.001 vs ACSF), or dynasore (85.7 ± 6.9%, n = 6, green, ***p ≤ 0.001 vs Gly, one-way ANOVA test). **(C)** Representative average NMDAR EPSC traces from mouse CA1 pyramidal neurons recorded at baseline and 20 minutes after bath-applied treatment of glycine (1 mM, 10 min, blue) or glycine plus dynasore (50 μM in intracellular solution, purple). **(D)**Histogram of averaged NMDAR EPSCs peak amplitude measured before (baseline) and after treatment of ACSF (98.9±4.1%, n = 6, grey), glycine (67.7 ± 3.6%, n = 15, blue, ***p ≤ 0.001 vs ACSF) and intracellularly applied dynasore (94.0 ± 4.7%, n = 6, green, ***p ≤ 0.001 vs Gly, one-way ANOVA test). **(E)** Scatter plots with representative traces showing the current (I)-voltage (V) relationship and reversal potential of NMDAR EPSCs before (white) and 25 min after glycine treatment (Gly 1 mM, 10 min, blue). Statistical significance (*p < 0.05) was detected at holding potential of −10 mV, +30 mV, +40 mV, +50 mV and +60 mV (n = 12, p < 0.05 between two plots, two-way ANOVA). (**F**) Histogram of averaged β2-adaptin/GluN1 ratio with glycine treatment (1mM, 10min) in hippocampal slices from wild-type mice (135.0 ± 8.10%, n = 13, ***p ≤ 0.001 vs ECS, Student t-test). Top, representative immunoblot of GluN1 and β2-adaptin with GluN1 immunoprecipitation.

Similarly, in hippocampal slices from wild-type C57Bl/6 mice, we observed that conditioning with bath-applied glycine produced a significant decrease in NMDAR EPSCs in CA1 pyramidal neurons (Fig. 3C,D). As in rat CA1 neurons, intracellular application of dynasore via the patch pipette prevented the glycine-primed decrease in NMDAR EPSCs (Fig. 3C,D). We compared the current-voltage (I-V) relationship and reversal potential of NMDAR-EPSCs at the beginning and the end of each recordings and found that glycine conditioning reduced the slope conductance but did not change the reversal potential (Fig. 3E). In addition, we found that glycine treatment led to a decrease of NMDAR EPSCs in neurons recorded continuously at holding potential of −10 mV (Supplementary Fig. 2). Thus, the glycine-primed depression did not depend up the neuron membrane potential being held at +60 mV, and this depression was not caused by a change of driving force for NMDARs. To test the possibility that conditioning with glycine might non-specifically depress excitatory synaptic transmission at Schaffer-CA1 pyramidal neuron synapses, we examined pharmacologically isolated AMPAR EPSCs (Supplementary Fig. 3). We found that conditioning with bath-applied glycine had no effect on the synaptic AMPAR responses and thus glycine did not cause depression of excitatory transmission non-specifically. We take these findings together as evidence that glycine primes a persistent, dynamin-dependent depression of NMDAR EPSCs at Schaffer-pyramidal neuron synapses in CA1 in rats and mice.

### Glycine stimulation drives association of AP2 with NMDARs from wild-type mouse hippocampal slices

As glycine primes a decline in synaptic NMDAR currents in hippocampal CA1 pyramidal neurons from rats and mice, we wondered whether glycine caused recruitment of AP2 to the NMDAR complex in the hippocampus. We used co-immunoprecipitation of AP2/NMDAR complexes from hippocampal slices from wild-type C57Bl/6 mice. Slices were conditioned with either control ECS or ECS + glycine in the presence of D-APV. We observed that glycine conditioning induced a significant increase in AP2/NMDAR association in mouse slices (Fig. 3F), which is consistent with our previous observation that glycine conditioning enhances the association of AP2 with NMDARs in rat hippocampal slices [24]. From our findings we conclude that glycine conditioning primes depression of synaptic NMDAR currents and enhances association of AP2 with the NMDAR complex for native receptors.

### N1 cassette of GluN1 prevents glycine-primed decline in synaptic NMDAR currents in hippocampal CA1 pyramidal neurons

In the adult rodent hippocampus *Grin1* mRNA is a mixture of transcripts that contain or that lack exon 5 [30, 31]. Thus, native NMDARs are predicted to be a mixture in which there are receptors with GluN1 subunits containing the N1 cassette and those with GluN1 lacking this cassette. That synaptic NMDARs are normally composed of a mixture of N1-containing and N1-lacking GluN1 is consistent with findings comparing wild-type mice with genetically-modified mice either constitutively lacking *Grin1* exon 5 (*Grin1*^Δ5^ termed GluN1a mice; Fig. 4A) or obligatorily expressing *Grin1* exon 5 (*Grin1*^Ω456^ termed GluN1b mice)[14]. To investigate the role of the N1 cassette of GluN1 in glycine-primed depression of synaptic NMDARs at Schaffer collateral synapses onto pyramidal neurons we compared the effects of glycine conditioning on NMDAR EPSCs in GluN1a versus GluN1b mice. Such a comparison is experimentally appropriate because basal synaptic transmission at these synapses in GluN1a mice is not different from basal transmission in GluN1b mice [14]. Here, we found that glycine conditioning led to a significant decrease in NMDAR EPSCs in CA1 pyramidal neurons in hippocampal slices from GluN1a mice (Fig. 4A,B). The proportionate depression of NMDAR EPSCs caused by conditioning glycine in GluN1a pyramidal neurons was not different from that of the glycine-primed depression in the wild type. In contrast to GluN1-1a, in CA1 pyramidal neurons in slices obtained from the GluN1b mice conditioning with bath-applied glycine failed to produce a change in the amplitude of NMDAR EPSCs (Fig. 4A,B).

**Figure 4.**
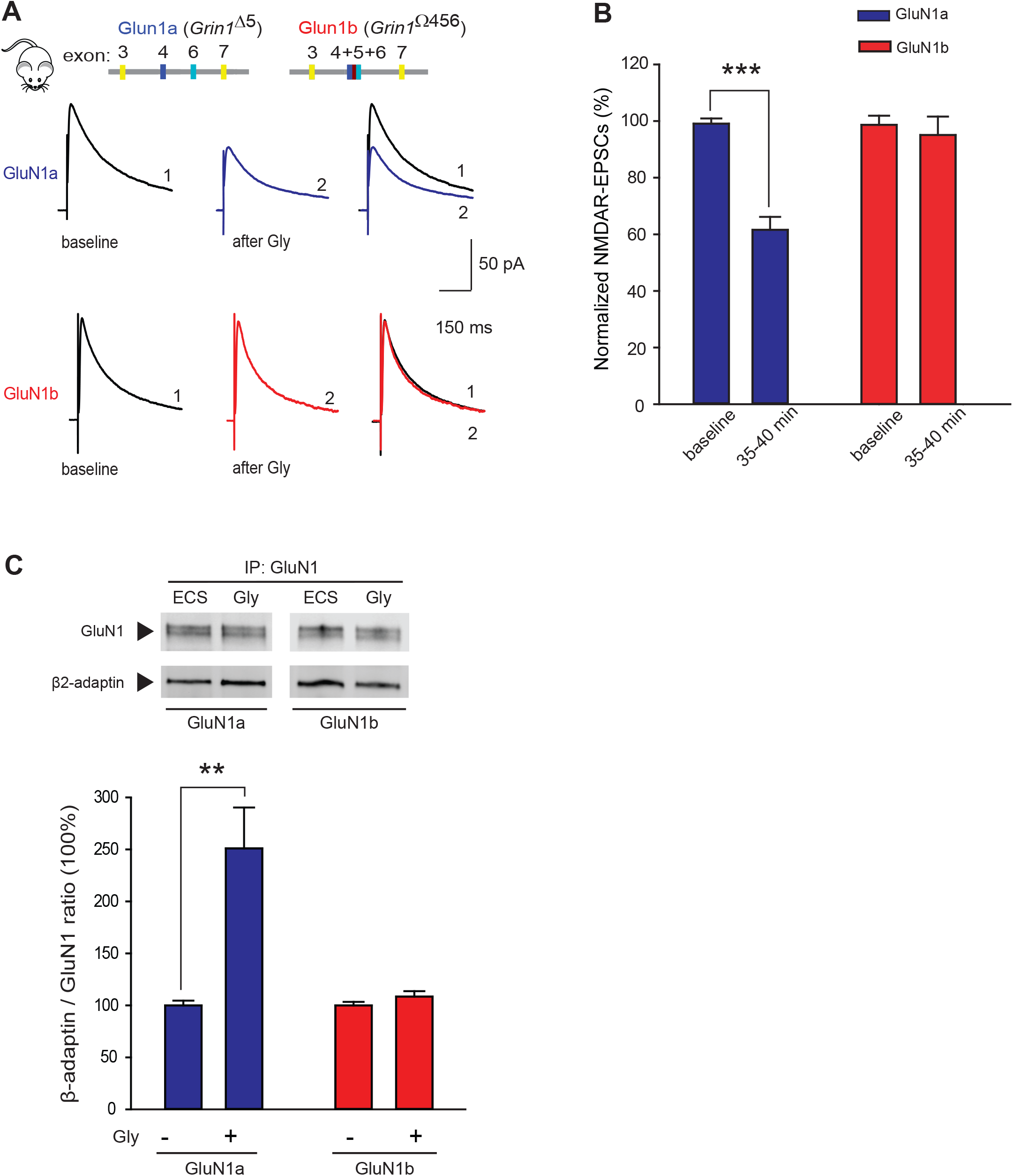
Glycine-primed decline in synaptic NMDAR current is dependent on the exclusion of the N1 cassette. **(A)** Top, schematic representation of *Grin1* loci for GluN1a and GluN1b mice depicting removal of exon 5 (*Grin1*^Δ5^) or fusion of exons 4 to 6 (*Grin1*^Ω456^). Bottom, representative average NMDAR EPSC traces from GluN1a (blue) and GluN1b (red) mice CA1 pyramidal neurons recorded at baseline (black) and 20 minutes after bath-applied treatment of glycine (1 mM, 10 min). **(B)**Histogram of averaged NMDAR EPSCs peak amplitude measured before glycine treatment as baseline and at the end of each recordings shown as: GluN1a (60.8 ± 4.8%, n = 13, ***p ≤ 0.001 vs baseline, blue), GluN1b (94.8 ± 4.2%, n = 7, p > 0.05 vs baseline, red). **(C)**Histogram of averaged β2-adaptin/GluN1 ratio with glycine treatment (1mM, 10min) in hippocampal slices from GluN1a (250.9 ± 43.0%, n = 6, **p < 0.01 vs ECS, blue) and GluN1b mice (108.4 ± 5.4%, n = 11, p > 0.05 vs ECS, red). Top, representative immunoblot of GluN1 and β2-adaptin with GluN1 immunoprecipitation. Student t-test is used for all statistic comparisons.

### N1 cassette prevents glycine-primed recruitment of AP2 to recombinant NMDARs

As forcing GluN1 subunits to contain the N1 cassette prevented glycine-primed depression of NMDAR EPSCs at Schaffer collateral synapses, we wondered whether the absence or presence of the N1 cassette of GluN1 affects glycine-stimulated recruitment of AP2 to the NMDAR complex in the hippocampus. We used co-immunoprecipitation of AP2/NMDAR complexes from hippocampal slices from GluN1a or GluN1b mice. Slices were conditioned with either control ECS or ECS + glycine. We observed that glycine conditioning induced a significant increase in AP2/NMDAR association in slices from GluN1a mice (Fig. 4C). But conditioning with glycine did not change AP2/NMDAR association was observed in slices from GluN1b mice (Fig. 4C).

From our findings we conclude that glycine conditioning primes depression of synaptic NMDAR currents and enhances association of AP2 with the NMDAR complex only for native receptors in which GluN1 subunits lack the N1 cassette. That is, expression of GluN1 subunits containing the N1 cassette prevents glycine-primed enhancement of AP2 association with NMDARs and also prevents depression of NMDAR ESPCs at Schaffer-collateral synapses of pyramidal cells.

### Glycine primed NMDAR internalization is absent in hippocampal interneurons

Schaffer collaterals make excitatory synapses onto inhibitory interneurons, as well as onto pyramidal cells in CA1. Thus, we tested whether glycine-primed depression of NMDAR EPSCs generalizes to other excitatory synapses in CA1 hippocampus. We made whole-cell patch-clamp recordings from interneurons dispersed within the stratum radiatum. We established that neurons recorded in stratum radiatum were interneurons using current clamp mode (Fig. 5A-C). These neurons showed a distinctive firing pattern and action potential peak amplitude in response to depolarizing current injection (Fig. 5A,B), and the amplitude of their afterhyperpolarization was much larger than that of pyramidal cells (Fig. 5C). To our surprise, we found that glycine conditioning had no effect on the amplitude of NMDAR EPSCs recorded from stratum radiatum interneurons in wild-type C57BL/6J mice (Fig. 5D,E). The lack of effect of glycine on interneurons contrasted with recordings from pyramidal neurons in these mice where we observed that conditioning with glycine treatment produced a gradual and sustained decrease in NMDAR EPSC amplitude (Fig. 5D,E), as expected from our findings above. Thus, glycine-primed depression of NMDAR EPSCs does not generalize to all excitatory synapses.

**Figure 5.**
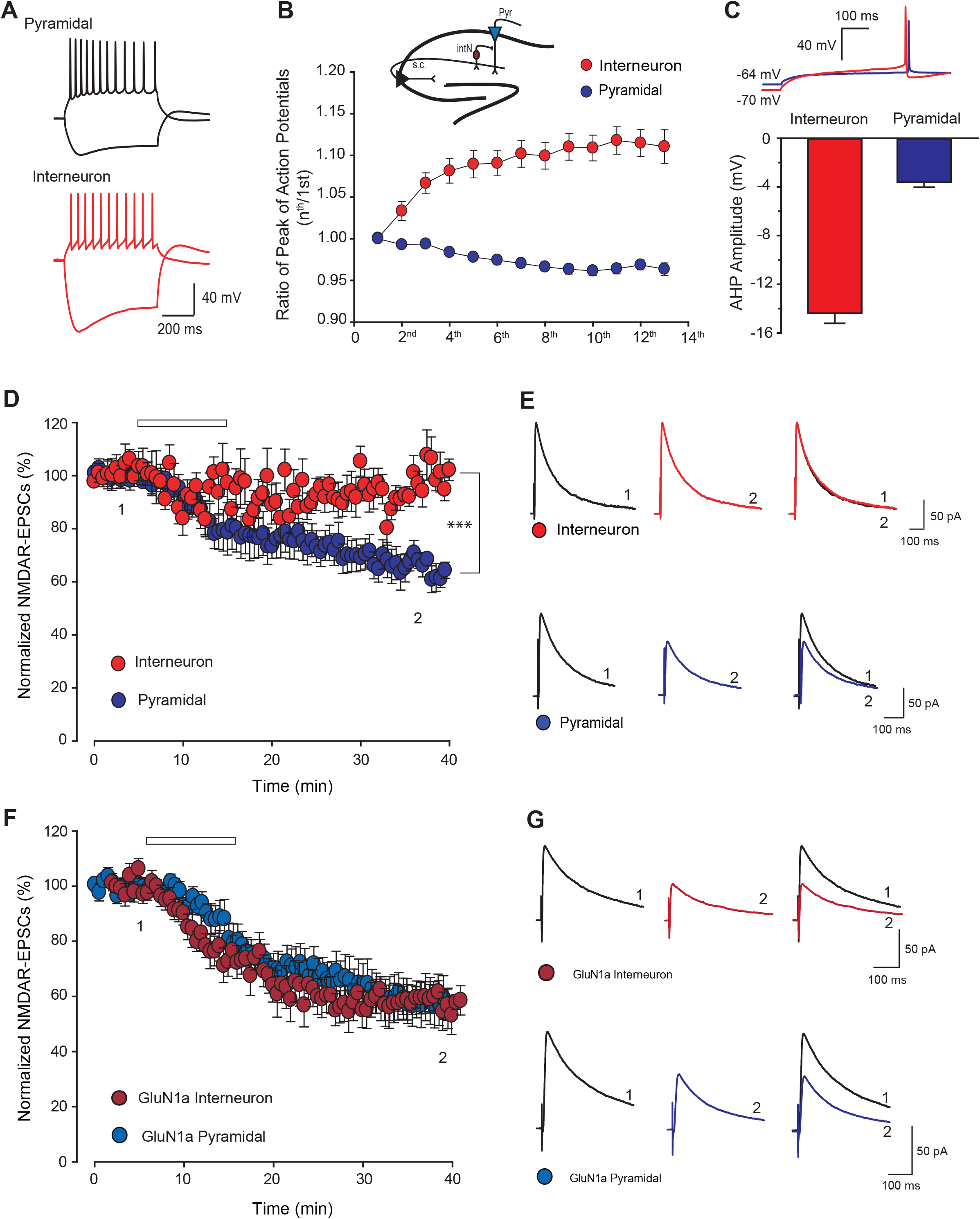
Glycine-primed NMDAR internalization is inhibited in hippocampal interneurons. **(A)** Representative traces of voltage responses to hyperpolarizing current (−200 pA, 600 ms) and action potentials to depolarizing current injections recorded in CA1 pyramidal neurons (+160 pA, 600 ms) and interneurons (+60 pA, 600 ms). **(B)** Plot of average action potential peaks in a train normalized to the first action potential for both cell types. Inset, diagram of a hippocampal slice showing recordings on CA1 pyramidal neuron (Pyr) and interneuron (intN) projected by Schaffer collaterals inputs. **(C)** Histogram of after-hyperpolarization potentials (AHPs) for both cell types. Top, representative traces of single action potential and AHPs evoked by rheobase depolarizations. **(D)** Scatter plot of NMDAR EPSC peak amplitude over time from mouse CA1 pyramidal (blue) and interneurons (red) recorded with treatment of 10 min bath-applied glycine (0.2 mM, white bar). The magnitudes of NMDAR EPSCs amplitude at time point 2 were as follows: Pyr (65.5 ± 3.9%, n = 7) versus intN (98.9 ± 5.4%, n = 6, ***p ≤ 0.001). Right, representative average NMDAR EPSC traces recorded at membrane potential of + 60 mV at the times indicated in (1 and 2). **(E)** Scatter plot of NMDAR EPSC peak amplitude over time from GluN1a mouse CA1 pyramidal (purple) and interneurons (dark red) recorded with treatment of 10 min bath-applied glycine (0.2 mM, open bar). The magnitudes of NMDAR EPSCs amplitude at time point 2 were as follows: Pyr (57.7 ± 7.8%, n =6) versus intN (57.8 ± 5.9%, n = 6, p > 0.05). Right, representative average NMDAR EPSC traces recorded at membrane potential of + 60 mV at the times indicated in (1 and 2). Student t-test is used for all statistic comparisons.

We considered the possibility that the lack of glycine-primed depression of NMDAR EPSCs was due to lack of GluN1a-containing NMDARs at the Schaffer collateral synapses onto inhibitory interneurons. As such, we predicted that forcing expression of only GluN1a-containing NMDARs at these synapses would recapitulate glycine-primed depression of the synaptic NMDAR currents. Consistent with this prediction, we found that glycine conditioning caused a use-dependent, progressive decline in NMDAR EPSCs recorded from stratum radiatum interneurons in hippocampal slices taken from GluN1a mice (Fig. 5F,G). Moreover, we found that neither the rate nor the extent of the glycine-primed decrease in NMDAR EPSCs in interneurons in GluN1a mice were different from those of the decrease in NMDAR EPSCs in pyramidal cells (Fig. 5F). Importantly, forcing expression of GluN1a-containing NMDARs did not alter the fundamental characteristics of these neurons – the firing pattern or after-hyperpolarization – that differentiate them from pyramidal neurons (Supplementary Fig. 4). Hence, we were clearly recording from interneurons in the slices from GluN1a mice. Thus, preventing expression of NMDARs with GluN1b, i.e. allowing only expression of GluN1a NMDARs, abrogated the lack of glycine-primed depression of NMDAR EPSCs in interneurons in CA1 hippocampus.

As a separate approach to examine the N1 status of NMDARs at the Schaffer collateral synapses onto inhibitory interneurons, we took the advantage of observations that recombinant NMDARs composed of N1 cassette-containing GluN1 exhibit faster deactivation rates than those without the N1 cassette [10, 11]. In wild-type mice we found that the decay times of the NMDAR EPSCs in pyramidal cells (230.3 ± 11.6, n = 10) was longer than that in interneurons (162.8 ± 13.7, n = 9, p < 0.05 t-test). In pyramidal cells, the decay times of NMDAR EPSCs were GluN1a > wild type > GluN1b (Supplementary Fig. 5B). By contrast, in interneurons the NMDAR-EPSC decay times were GluN1a >> wild type ~ GluN1b (Supplementary Fig. 5D). We interpret these findings as evidence supporting the conclusion that in interneurons in wild-type mice synaptic NMDARs are composed predominantly, if not exclusively, of N1-containing GluN1subunits. Taking our findings together, we conclude that the lack of glycine-primed depression of NMDAR EPSCs in inhibitory interneurons in the wild-type is due to lack of GluN1a-containing NMDARs at Schaffer collateral synapses on these interneurons.

## Discussion

Here we find that alternative splicing of GluN1 differentially gates glycine-primed internalization of NMDARs. Using heterologously-expressed recombinant NMDARs, we observed that receptors lacking the N1 cassette in GluN1 show glycine-stimulated recruitment of AP2, glycine-primed NMDAR internalization and glycine-primed depression of NMDAR currents. In stark contrast, we discovered that recombinant NMDARs with GluN1 containing the N1 cassette failed to show any of these three signature features of non-ionotropic NMDAR signalling caused by glycine. The lack of glycine stimulated non-ionotropic signalling was observed regardless of which of the C-terminal cassettes (C1, C2, C2’) of GluN1 was present. That is to say, the permissive effect of the lack of N1 was observed regardless of the C-terminal cassette. Moreover, the blockade glycine priming by the N1 cassette – the permissive effect of its lack – was found with recombinant NMDARs containing either GluN2A or GluN2B. Thus, the splicing of exon 5, but splicing of other GluN1 exons, uniquely controls priming by glycine, and the effects of the presence, or lack, of exon 5 occur regardless of which GluN2 subunit the receptor is composed.

As glycine priming is initiated by binding of glycine to the receptor[28], it is conceivable that the lack of this glycine-induced non-ionotropic signalling by NMDARs with the N1 cassette in GluN1 might be due simply to a lack of glycine efficacy on such receptors. We previously identified a single amino acid substitution, A714L, in the ligand binding domain of GluN1 that likewise prevents glycine-priming[25]. The A714L mutation dramatically reduces the potency of glycine to induce channel gating and this reduce binding efficacy is a simple explanation as the cause of the lack of the ability of that mutant receptor to undergo glycine priming. However, here, neither the potency nor efficacy of glycine for its ionotropic effect, i.e. channel gating when glycine is bound to the receptor together with glutamate, are compromised in receptors containing the N1 cassette [11, 14]. Thus, a simple lack of glycine binding or ability of glycine binding to stabilize an altered conformation of the receptor cannot be the mechanism for the blockade of priming by the N1 cassette. As a corollary, reduction of glycine binding is not a prerequisite to block non-ionotropic signalling by GluN1.

We have proposed a schema for glycine priming of NMDARs whereby binding of glycine to its cognate site in GluN1, in the absence of binding of glutamate to GluN2, is coupled to conformation changes in the receptor complex initiated in the extracellular region that are transmitted across the plasma membrane where, intracellularly, these changes allow the binding of the AP2 complex [28]. As a consequence, the AP2 bound state of the NMDAR is then able to undergo dynamin-dependent endocytosis upon the ligation of the receptor glutamate, and glycine. As we presently find that glycine does not recruit AP2 to NMDARs composed of N1-containing GluN1, we hypothesize that the N1 cassette prevents, or does not permit, the conformation rearrangements required to couple the binding glycine extracellularly to the recruitment of AP2 intracellularly. This effect of the N1 cassette could come about in a number broad conceptual ways: i) by N1 sterically preventing the receptor from taking on the conformation, or by destabilizing this conformation, that links binding of glycine to transmembrane signalling, ii) by N1-containing receptors still signalling across the membrane, but through conformational rearrangements that prevent, or don’t permit, the intracellular binding of AP2, iii) by the presence of N1 changing extracellular or intracellular post-translational modifications in GluN1, or GluN2 subunits, that are necessary for transmembrane signalling or recruitment of AP2, or iv) by the presence of N1 causing changes in the composition of accessory proteins, lipids, or other molecules, in the NMDAR complex associated proteins such that a molecule necessary for transmembrane signalling or recruitment of AP2 is missing, or a molecule blocking these steps is recruited. Cryo-electron microscopy structural data shown that the 21-amino acid residues of the N1 cassette create an interface area between the amino-terminal domain and the glycine ligand-binding domain [32]. Such an interface may support any of these four broad ways in which the N1 cassette could prevent glycine priming. Hence, our present findings open up a number of mechanistic possibilities to be addressed in the future.

Glycine-primed depression of NMDAR currents neurons had only been previously shown for NMDAR-mediated responses evoked by exogenous agonists in acutely isolated hippocampal neurons or in spontaneous EPSCs in cells in culture[24]. Here we demonstrate that NMDAR EPSCs at Schaffer collateral synapses in CA1 pyramidal neurons in acute hippocampal slices from wild-type mice are primed by glycine for subsequent dynamin-dependent depression when the synaptic receptors are activated. We also demonstrate that glycine treatment causes recruitment of AP2 to the NMDAR complex in mouse hippocampal slices. Thus, our present findings extend previous observations on the effects of glycine on native NMDARs and indicate that glycine priming occurs at NMDARs at synapses *ex vivo*.

In contrast to our findings in the wild type, in slices from GluN1b mice NMDAR EPSCs at Schaffer collateral synapses in pyramidal neurons did not show glycine-primed depression. Nor was AP2 recruited to the NMDAR complex by glycine in GluN1b mice. On the other hand, in GluN1a mice there was robust glycine-primed depression of NMDAR EPSCs at pyramidal neuron Schaffer collateral synapses, and glycine-primed recruitment of AP2. Thus, as predicted from our findings with heterologously expressed recombinant NMDARs, forcing inclusion of the N1 cassette in native NMDARs prevents glycine priming. That is, splicing out exon 5 in pyramidal neurons permits this non-ionotropic signalling by glycine.

Surprisingly, given the robust and highly consistent glycine-primed depression of NMDAR EPSCs at Schaffer collateral synapses onto pyramidal neurons, at Schaffer collateral synapses onto interneurons in stratum radiatum in wild-type mice glycine treatment had no effect on NMDAR ESPCs. Glycine-primed depression of NMDAR EPSCs in interneurons was recapitulated in slices from GluN1a mice indicating that preventing inclusion of exon 5, and the N1 cassette, is sufficient to allow non-ionotropic signalling by glycine in these neurons. The lack of glycine-primed depression of NMDAR EPSCs and the faster decay time of NMDAR EPSCs in interneurons in the wild type, and the rescue of glycine priming in GluN1a mice, together suggest that synaptic NMDARs in interneurons are normally composed of GluN1 subunits containing the N1 cassette, that is exon 5 is normally included in GluN1 mRNA in these neurons. This interpretation is supported by recent evidence from FACS sorted cortical neurons in which interneurons have much lower levels of GluN1 transcripts lacking exon 5 than do excitatory neurons[30]. Amongst the major subtypes of interneurons (parvalbumin, PV; somatostatin, SST; vasoactive intestinal polypeptide, VIP) the lowest levels transcripts lacking exon 5, on average about 10%, are expressed in the PV interneurons. PV interneurons are prominent in stratum radiatum and the very low level of exon 5-lacking transcripts in these cells is consistent with our findings of no detectable glycine-primed depression of synaptic NMDAR currents in the interneurons we recorded.

In pyramidal cells, the level of depression of NMDAR EPSCs primed by glycine, approximately 40% depression, was similar to the level with recombinant receptors expressing GluN1-1a (cf. Fig 2A,B with Fig 3A,B). In GluN1a mice, the degree of glycine-primed depression of NMDAR EPSCs in pyramidal neurons was not greater than that in the wild type (cf. Fig 3A,B with Fig 4A,B). Thus, in pyramidal neurons in wild-type mice the non-ionotropic glycine signalling effect of GluN1 subunits lacking the N1 cassette appears to be maximal and dominates over any effect of subunits containing N1 that may be present at synapses[14]. This interpretation is supported by the finding that in excitatory cortical neurons greater than 90% of GluN1 mRNA transcripts have been found to lack exon 5 [30]. The differential level of exon 5 inclusion in excitatory vs inhibitory neurons – with nearly all transcripts in the former lacking exon 5 and the majority in the latter containing exon 5 – provides an explanation for the dramatic increase in glycine-stimulated recruitment of AP2 to the NMDAR complex in hippocampal extracts from GluN1a vs wild-type mice (cf. Fig 3G and Fig 4D). That is, forcing the exclusion of exon 5, and thereby the N1 cassette, in interneurons of GluN1a mice allows glycine-primed recruitment of AP2 to NMDARs in these neurons, which was absent in the wild type. Whereas in pyramidal neurons such recruitment is near maximum even in wild-type mice and neither AP2 recruitment nor glycine-primed depression of NMDAR EPSCs can be increased in these neurons in GluN1a mice.

That glycine-stimulated depression of NMDAR EPSCs can be prevented in pyramidal neurons, in GluN1b mice, and imposed on interneurons, in GluN1a mice, demonstrates cell-type specific functional consequences attributable to the differences in splicing of exon 5. This differential alternative splicing in pyramidal neurons vs inhibitory interneurons may be mediated through differential action of splicing factor(s) in these two cells types. An example of one such splicing factor is the RNA-binding protein NOVA2 which has been found to control unique RNA splicing programs in inhibitory and excitatory neurons [33]. The identity, or identities, of the splicing factor(s) responsible for the differential splicing of GluN1 exon 5 remains to be determined.

Additional examples of non-ionotropic signalling of NMDARs stimulated by glycine are increasingly reported [34, 35]. Our present findings raise the possibility that these non-ionotropic glycine signalling events might also be gated by exon 5 splicing, and the presence or absence of the N1 cassette in GluN1. Differential splicing of *Grin1* exon 5 has been found to control the level of hippocampal LTP at Schaffer collateral synapses on pyramidal neurons[14]. The reduced LTP observed in GluN1b mice, could not be explained by a number of factors, including basal transmission, synaptic NMDAR or AMPAR currents, NMDAR to AMPAR ratios, magnesium block potency, nor allosteric potentiation by magnesium or spermine[14]. As a result, it is conceivable that non-ionotropic signalling by glycine through GluN1 may be responsible for these differences, and that the lack of the N1 cassette may permit transmembrane signal transduction that is necessary for LTP in hippocampal pyramidal neurons.

Our findings altogether uncover a previously unanticipated molecular function for the N1 cassette, encoded by exon 5, in controlling non-ionotropic glycine signalling through the GluN1 subunit of NMDARs. We suggest that differential alternative splicing of exon 5 may play important, cell-specific signalling roles as demonstrated by the distinct phenotypes between CA1 neurons and interneurons within the hippocampus. Our findings raise the possibility that the difference in exon 5 splicing in excitatory vs inhibitory neurons generalizes throughout the CNS and may thereby have widespread consequences for NMDAR-dependent integration and circuit function.

## Acknowledgements

We thank technical assistance provided by Vivian Wang and Janice Hicks. This study was supported by grants from CIHR to M.W.S. (FDN-154336) and a Restracomp post-doctoral fellowship to V.R.

## Author contributions

The authors declare no competing financial interests. H.L., V.R., D.C., L.H., and J.E.C. performed the experiments for this work. H.L, A.S.S. and M.W.S. designed the research. H.L., V.R., A.S.S. and M.W.S. wrote the manuscript. M.W.S. supervised the project.

## Methods

### Animal Usage

For experiments, GluN1a and GluN1b heterozygous mice were bred to generate GluN1a and GluN1b homozygous mice and their respective WT littermates. Mice from both GluN1a and GluN1b colonies were housed in the same room and in the same racks. The WT littermates of GluN1b homozygotes did not differ from the WT littermates of GluN1a homozygotes[14]. Experimental use of animals was in accordance with policies of the Hospital for Sick Children Animal Care Committee and the Canadian Council on Animal Care.

### Cell culture and transfection

Human embryonic kidney cell line (HEK293) cells (3 × 10^4^ cells per cm^2^) were plated onto 6-well culture dishes coated with poly-D-lysine. HEK293 cells were cultured with Dulbecco’s Modified Eagles Media (DMEM) (Invitrogen) supplemented with 10% fetal bovine serum (Invitrogen) and 1% penicillin-streptomycin (Wisent); 37°C, 5% CO_2_. For electrophysiological recordings, HEK293 cells were seeded on poly-D-lysine coated glass coverslips at 1.0 × 10^5^ cells per well in a 6-well dish 24 hrs before transfection. FuGene HD (Promega BioSciences) was used for all transfections. Transfections contained expression plasmids of GluN1, GluN2 (either GluN2A or GluN2B); and PSD-95 at a ratio of 1:4:0.5 [25]. For electrophysiological recordings an enhanced green fluorescent protein (eGFP) plasmid was included in the transfection at a ratio of 0.5 with respect to GluN1. After transfection, cells were maintained in DMEM supplemented with 10% fetal bovine serum and D-APV (500 μM; Tocris) for 48 hrs before experiments.

### Colorimetric cell enzyme-linked immunosorbent assay (ELISA)

HEK293 cells were seeded at 2.5 × 10^5^ cells per well in 12-well plates 24 hrs prior to transfection and grown for 48 hrs after transfection. The media was replaced with cold ECS (in mM: NaCl 140, CaCl_2_ 1.3, KCl 5, HEPES 25, and glucose 33) and plates were maintained on ice to inhibit membrane trafficking. To label cell surface NMDARs, cells were incubated for 1 hr at 4°C with extracellular domain anti-GluN1 antibody (BD biosciences; 2 μg ml^−1^) followed by two washes with ice cold phosphate buffered saline (PBS, Wisent). Cells were treated with ECS containing 100 μM glycine and 100 μM D-APV at 37°C for 5 min or with ECS lacking glycine for control condition. Solution was then replaced with ECS containing 50 μM NMDA plus glycine 1 μM for at 37°C 10 min. Cells were fixed with cold 4% paraformaldehyde in PBS for 10 min on ice and washed twice with ice cold PBS. Cells were incubated for 1 hr at room temperature with a horseradish peroxidase conjugated secondary antibody (1:1000, Amersham) diluted in PBS. ELISA reaction procedure was followed as described by manufacturer (OPD; Sigma). The optical density of the supernatant was read on a spectrophotometer at 492 nm. The levels of cell surface expression of NMDARs were presented as a ratio of colorimetric readings measured on cells not subject to the OPD reagent.

### Confocal microscopy of NMDAR internalization

GluN1 subunit was modified to include a 13 amino acid BBS sequence at the N-terminus, immediately following the signal sequence, to detect internalization by confocal microscopy as previously described [25].

Surface BBS-NMDARs were labeled with 3 μg ml^−1^ BTX-CypHer5E at 4°C for 30 min, washed twice with cold PBS and treated with control ECS or ECS with 100 μM glycine and 100 μM D-APV for 5 min at 37°C. The labeling was not sufficient to saturate all the BBS-NMDARs. Live cells were then treated with control ECS or NMDA (50 μM) plus glycine (1 μM) for 10 min. After washing with cold ECS, cells were incubated with BTX-AF488 (Molecular Probes; 10 mM) at 18°C for 20 min. Cells were washed to remove unbound BTX-AF488 and then imaged using confocal microscopy (Olympus IX81). Expression of the BBS-tagged receptors was verified by extracellular acidification with acetate buffer (pH 5.5) to mimic luminal endocytic vesicle pH. Images were captured using a Hamamatsu BackThinned EM-CCD camera and Volocity software (Perkin Elmer).

### Immunoprecipitation and immunoblot for β2-adaptin with NMDARs

For hippocampal immunoprecipitates, hippocampal slices were prepared from adult mice (C57Bl/5, Jackson Laboratory) same as described for electrophysiology recording. Six slices (400 μm thick) were treated with D-APV (100 μM) with or without glycine (1 mM) for 10 min in ECS containing strychnine (1 μM), bicuculline (20 μM) and tetrodotoxin (1 μM). For recombinant NMDAR containing immunoprecipitates, transfected HEK293 cells expressing NMDARs were treated for 5 min with glycine (100 μM) plus D-APV (100 μM) or control ECS alone (with D-APV).

Slices or cells were homogenized in ice cold lysis buffer: 50 mM Tris-HCl (pH 8.0), 150 mM NaCl, 2 mM EDTA, 0.1% SDS, 1% NP-40, 0.5% sodium deoxycholate, protease inhibitor cocktail tablet (Roche) and phosphatase inhibitor cocktail set III (Millipore-Sigma). Insoluble material was removed by centrifugation at 14,000g for 20 min at 4°C. Soluble proteins were incubated overnight with 2.5 μg of anti-GluN1 antibody (BD Pharmingen). Immune complexes were isolated by the addition of 20 μL of protein G– Sepharose beads (GE Healthcare) followed by incubation for 2 hr at 4°C. Immunoprecipitates were washed three times with lysis buffer, resuspended in laemmli sample buffer, and boiled for 30 min at 70°C. The samples were subjected to SDS– polyacrylamide gel electrophoresis (PAGE) on a 4–20% gradient gel and transferred to a polyvinylidene difluoride (PVDF) membrane. Membranes were immunoblotted with polyclonal anti-GluN1 (1:1000, Genetex) and anti-β2-adaptin (1:500, BD Biosciences) overnight and then probed with their respective secondary antibodies (Jackson Immunoresearch). Blots were imaged using Odyssey imagining system and quantified with Image Studio Lite (LI-COR Biosciences. Serial dilutions were used to confirm that under these experimental conditions signal intensities for GluN1 or adaptin β2 were linear over a 50-fold range. We note that immunoprecipitating with a non-specific IgG caused no detectable precipitation of GluN1 or adaptin β2.

### Whole-cell recording in recombinant NMDARs

Whole-cell recordings were made at room temperature (20–22°C). After formation of a whole-cell configuration, the recorded neurons were voltage-clamped (−60 mV) and lifted into the stream of solution supplied by a computer controlled, multi-barrelled fast perfusion system. The extracellular solution was composed of (in mM): NaCl 140, KCl 5.4, CaCl_2_ 1.3, HEPES 25 and D-glucose 33, Glycine 0.001 (pH 7.4, osmolarity 330 mosM). The intracellular solution contained (in mM): CsF 140, BAPTA 10, Hepes 10 and MgATP 2 (pH 7.2 with CsOH and osmolarity 300 mosM). NMDAR currents were evoked by short applications (3 s duration) of NMDA (50 μM) and glycine (1 μM) every 60 sec. Glycine priming experiments was made by a 5 min application of glycine (100 μM plus D-APV 100 μM) (D-APV was included to avoid activating NMDAR channel gating). After the 5 min application, the glycine priming solution was washed for 1 min with control ECS, before re-probing NMDAR activity with the test NMDA and glycine applications every 60 sec. NMDAR currents were recorded using an Axopatch 1-D amplifier, data were digitized with DigiData1200A, filtered (2 kHz), digitized at 10 kHz and acquired by the pClamp9.0 software (Molecular Devices). Recordings in which the series resistance varied by more than 10% were rejected. NMDA-evoked current data are presented as percentage of the peak mean current (I) normalized to the initial response (I_0_). All data are presented as means ± s.e.m. Once whole-cell configuration was achieved, we allowed 10–15 min for diffusion to the cell cytoplasm and then started recording NMDA-evoked currents.

### Hippocampal slice electrophysiology

We prepared parasagittal hippocampal slices (300 μm) in ice-cold ACSF from anesthetized (20% wt vol^−1^) urethane, intraperitoneal (i.p.) mice (C57BL/6) (22-28 days) or Sprague Dawley rats (15-21 days) and placed them in a holding chamber (30°C) for 40 min and then allowed it to passively cool down to room temperature (21 to 22 °C for ≥ 30 min) before recording. We then transferred a single slice to a recording chamber and superfused with ACSF at 4 ml min^−1^ composed of (in mM) 124 NaCl, 2.5 KCl, 1.25 NaH_2_PO_4_, 2 MgCl_2_, 11 D-glucose, 26 NaHCO_3_ and 2 CaCl_2_ (pH 7.40, osmolality 305 mOsm) saturated with 95% O_2_ (balance 5% CO_2_) at room temperature, pH 7.40, osmolality 310 mOsm. We evoked synaptic responses by stimulating Schaffer collateral afferents using bipolar tungsten electrodes located ~50 μm from the pyramidal cell body layer in CA1. Whole-cell patch-clamp recordings of CA1 pyramidal neurons were carried out using the visualized method (Zeiss Axioskop 2FS microscope). For mice slice recording, patch pipettes (4–5 MΩ) containing (in mM): Cs gluconate 117, CsCl 10, BAPTA 10, CaCl_2_ 1, HEPES 10, ATP-Mg 1, QX-314 10, GTP 0.3 (pH 7.25, osmolality 290 mOsm). Pipette solution for voltage-clamp recordings from rat slices contains (in mM): Cs gluconate 132.5, CsCl 17.5, EGTA 0.2, HEPES 10, ATP-Mg 2, QX-314 5, GTP 0.3 (pH 7.25, osmolality 290 mOsm). Hippocampal interneurons were identified by their location in stratum radiatum of CA1 region and their electrophysiological characteristics. Action potentials were evoked by a serial of current injections with patch pipettes containing (in mM): K gluconate 122.5, KCl 17.5, EGTA 0.2, HEPES 10, ATP-Mg 4, Na phosphocreatinine 10, GTP 0.3 (pH 7.25, osmolality 290 mOsm). Testing stimuli (0.1 ms in duration) were delivered at a frequency of 0.1Hz to evoke synaptic transmission. NMDAR-mediated EPSCs were pharmacological isolated by blockade of AMPA receptors with bath-applied CNQX (10 μM); bicuculline (10 μM) was included in the bath to block GABA_A_-receptor-mediated transmission; Glycine priming treatment (Gly 1 mM, 10 min) was bath applied after stable baseline recording. Strychnine (10 μM) was also included throughout the experiments to prevent the activation of glycine receptor during the glycine priming treatment. We amplified raw data using a MultiClamp 700B amplifier and a Digidata 1322A acquisition system sampled at 10 KHz and analyzed the data with Clampfit 10.6 (Axon Instruments).

### Quantification and statistical analysis

All experiments were performed with at least 3 individual mice or cell cultures. Number of samples/individuals used in each experiment are reported in the respective figure legends. All plots were composed in R or SigmaPlot. Sigmaplot or R were used for statistically analysis. All data were first determined to be normal or not before the appropriate parametric or non-parametric test were applied. Specific tests used to assess significance are mentioned for each experiment in their respective figure legends. Significance are based on an alpha level of 0.05 throughout this study. Values are mean ± SEM. with statistical significance indicated with *p < 0.05, **p ≤ 0.01 and ***p ≤0.001.

### Data availability

Data supporting the findings of this study are available within the article and its Supplementary Information files and from the corresponding author on reasonable request.

---

**Suppl. Figure 1.**
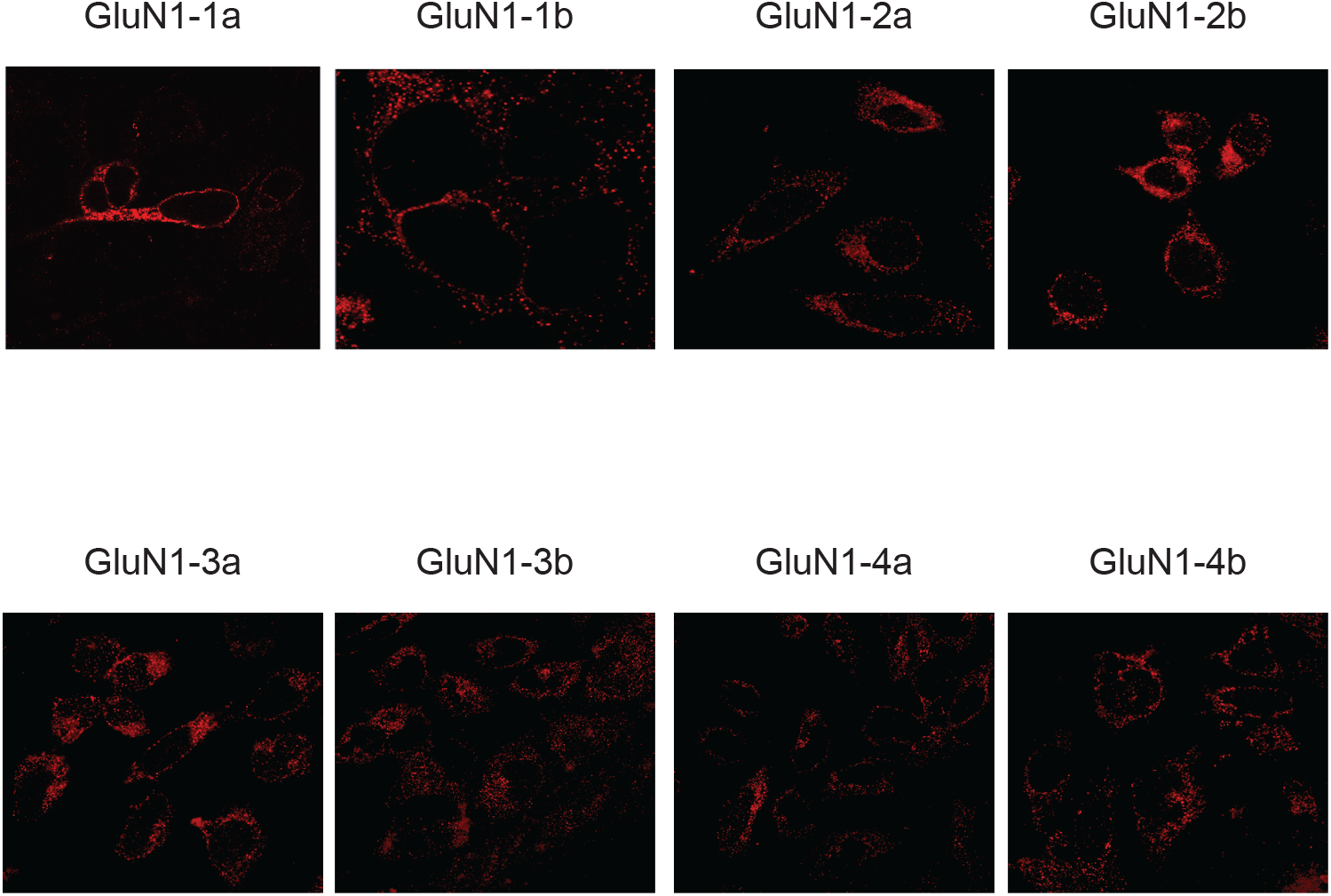
GluN1 splice variants expression in HEK293 cells. Confocal images of non-permeabilized HEK293 cells expressing GluN1 isoforms co-expressed with GluN2A. Surface receptors are labelled with anti-GluN1 antibody.

**Suppl. Figure 2.**
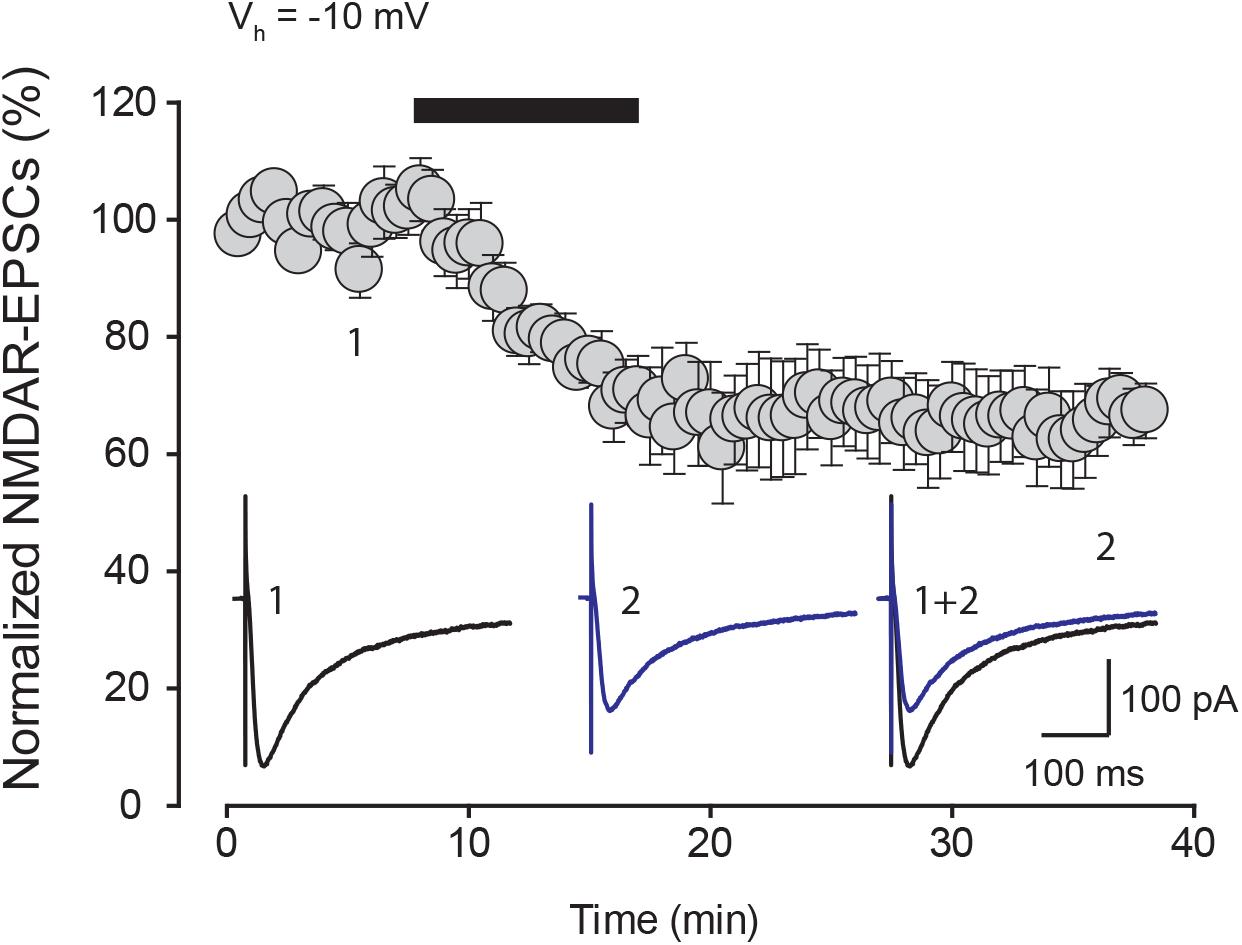
Glycine-primed decrease of synaptic NMDAR EPSCs recorded at holding potential of −10mV. Time series of normalized NMDAR EPSC peak amplitude from mouse CA1 pyramidal neurons recorded at holding potential of −10 mV. Slices were conditioned by bath-applied glycine (1 mM, 10 min, black bar). The magnitude of NMDAR-EPSCs 20 min after glycine was (65.5 ± 6.7%, n = 5, p ≤ 0.001 vs baseline, Student t-test). bottom, representative average NMDAR EPSC traces recorded at the times indicated (1 and 2).

**Suppl. Figure 3.**
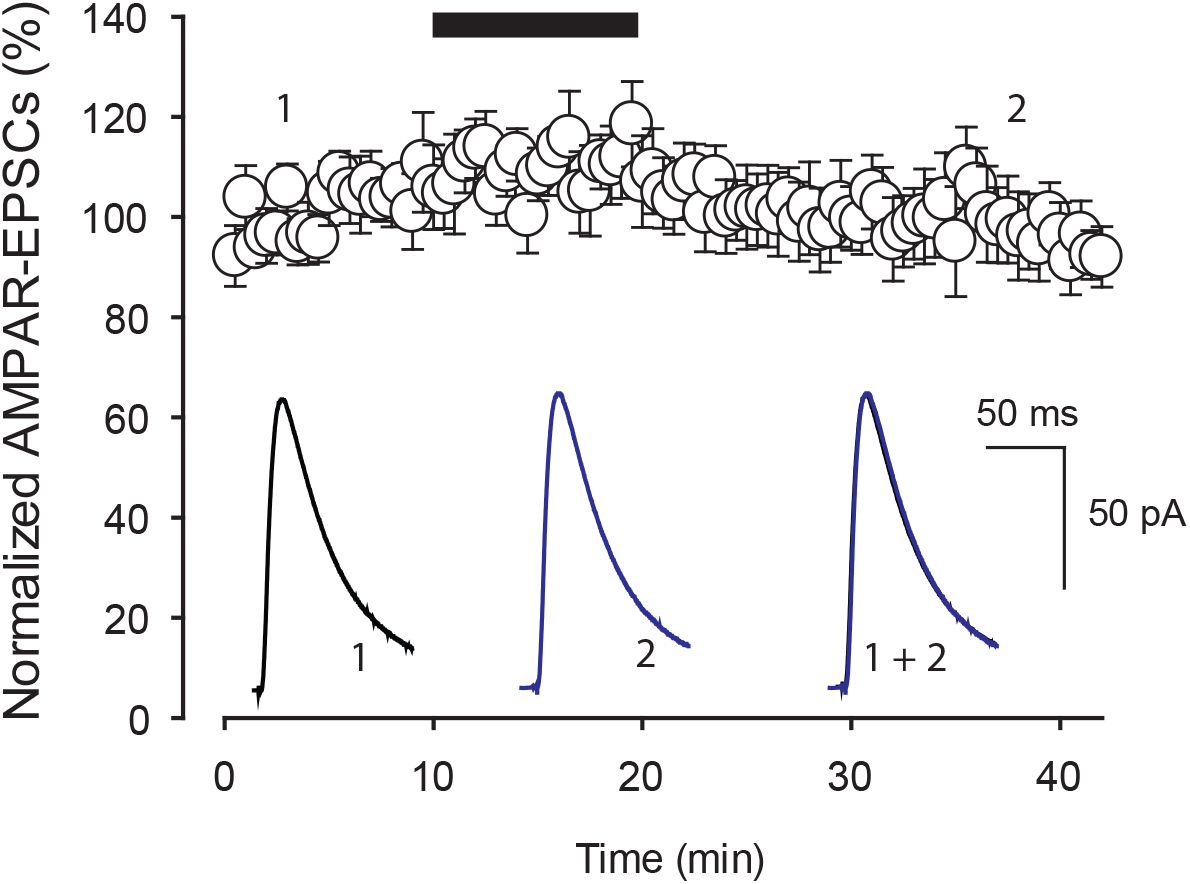
Glycine does not affect hippocampal synaptic AMPAR-EPSCs. Time series of normalized synaptic AMPAR-EPSC peak amplitude recorded from hippocampal CA1 pyramidal neurons. Glycine (1 mM) was applied for 10 min as indicated by the black bar. The magnitude of AMPAR-EPSCs 20 min after glycine was (95.7% ± 6.9%, n = 5, p > 0.05 vs baseline, Student t-test). Top representative traces of pharmacologically isolated AMPAR EPSCs recorded at + 60 mV right before and 20 min after glycine treatment. Top representative average AMPAR EPSC traces recorded at the times indicated (1 and 2). Each trace is an average of 10 consecutive traces.

**Suppl. Figure 4.**
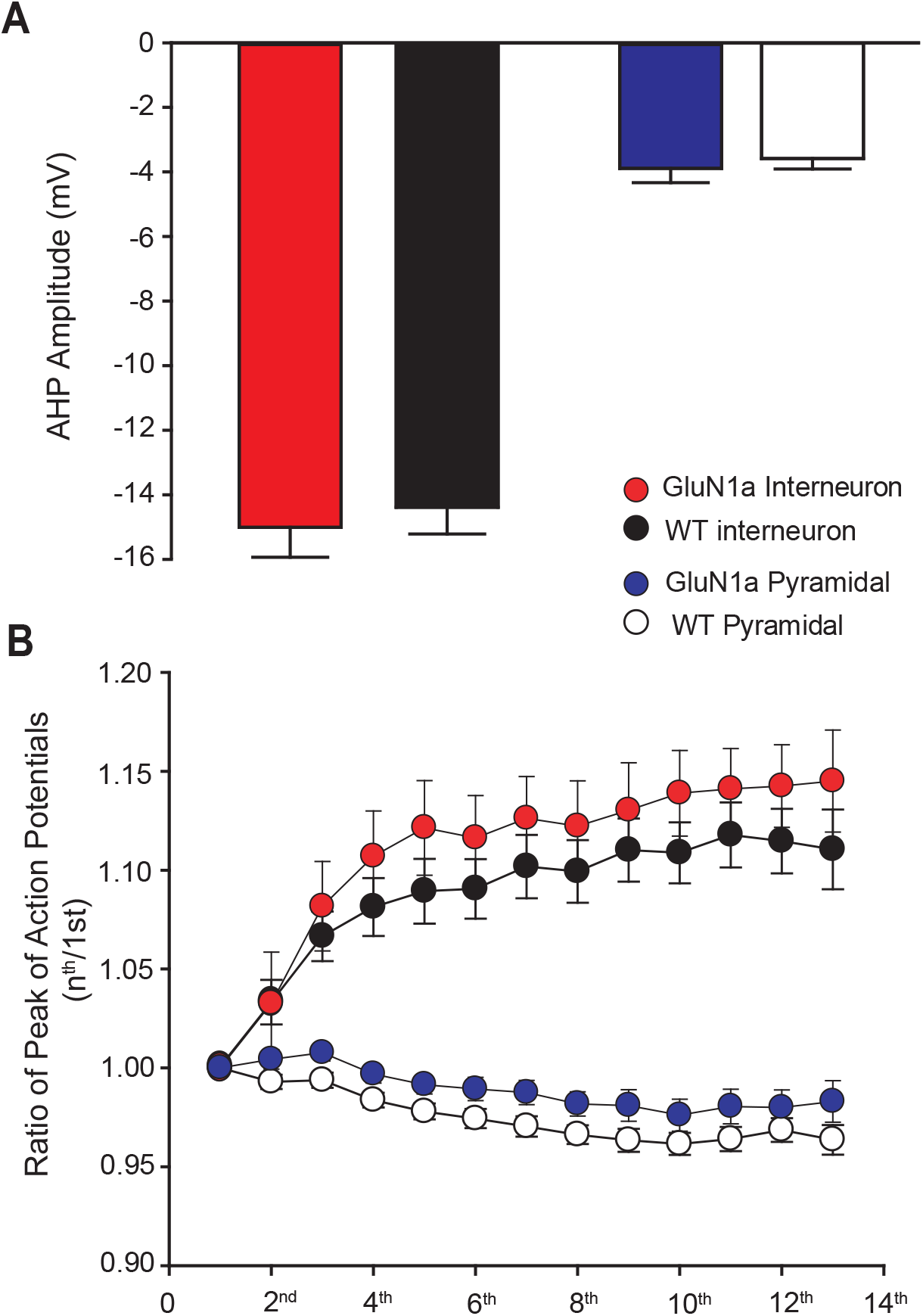
Electrophysiological characterizations of pyramidal and interneurons from GluN1a mice. **(A)** Histogram of after-hyperpolarization potential (AHPs) for GluN1a interneurons (red) and GluN1a pyramidal neurons (blue) compared to wild type interneurons (black) and wild type pyramidal neurons (white). Single action potential and AHPs evoked by rheobase depolarizations. **(B)** Plot of average action potential peaks in a train normalized to the first action potential for GluN1a interneurons (red), GluN1a pyramidal neurons (blue), wild type interneurons (black) and wild type pyramidal neurons (white).

**Suppl. Figure 5.**
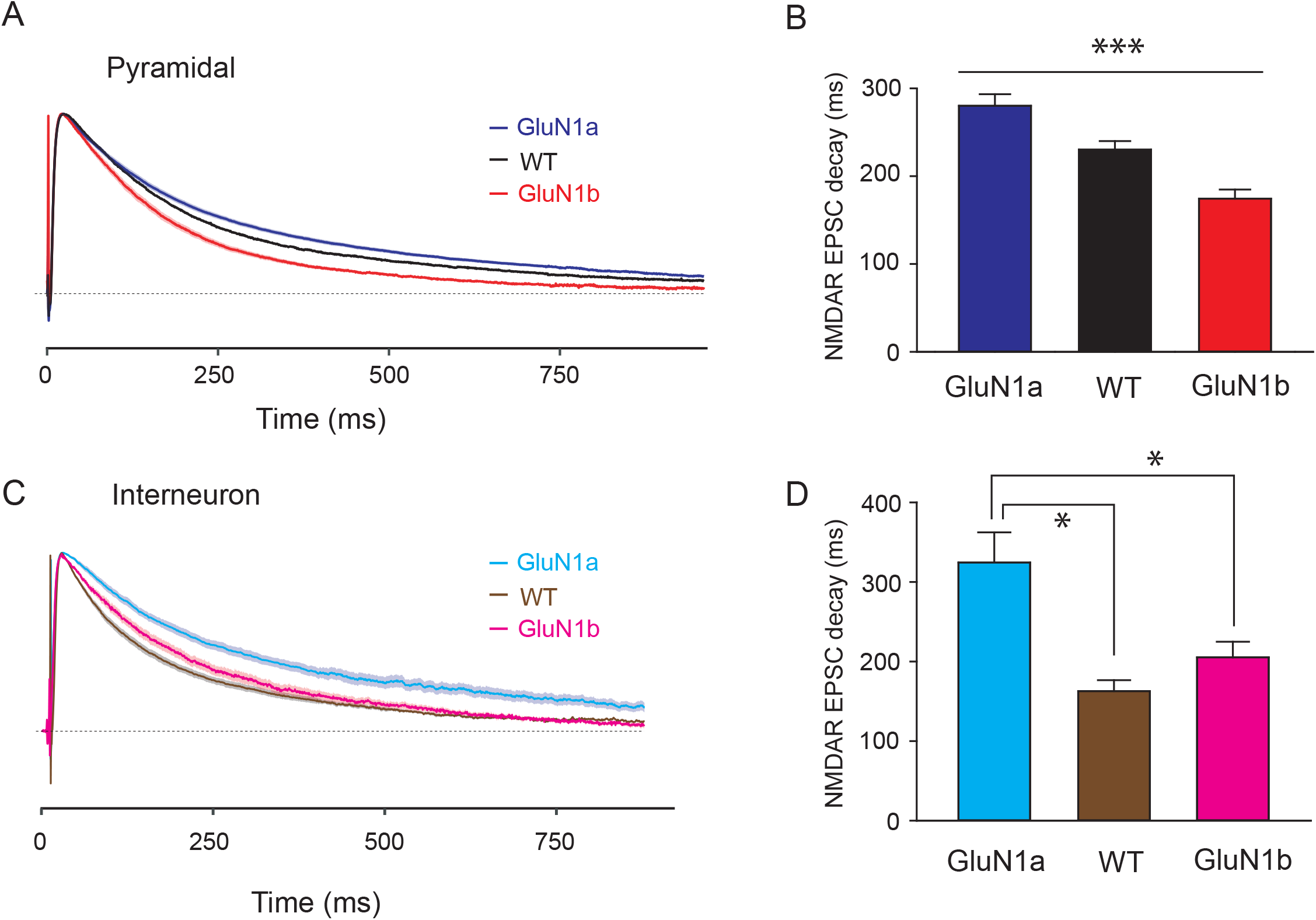
N1 cassette affects the decay time of NMDAR-EPSCs in CA1 pyramidal and interneurons. **(A)** Average normalized NMDAR-EPSC traces at + 60 mV recorded in CA1 pyramidal neurons from GluN1a (n=9), GluN1b (n=18) and wild-type (n=10) mice. **(B)** Histogram of the decay time of NMDAR-EPSCs based on the data shown in (A). There is a statistically significant difference among GluN1a, wild-type and GluN1b (p = < 0.001 One-way ANOVA test). **(C)** Average normalized NMDAR-EPSC traces at + 60 mV recorded in CA1 interneurons from GluN1a (n = 12), GluN1b (n = 6) and wild-type (n = 9) mice. **(D**) Histogram of the decay time of NMDAR-EPSCs based on the data shown in (C). There is no statistic significant difference between wild-type (n = 9) and GluN1b mice (n = 6) in interneurons (p > 0.05 One-way ANOVA test). The decay constant was calculated by the time (ms) it takes for the trace to decrease to 1 / e ≈ 36.8% from its peak. All traces are presented as mean (darker line) with standard error (lighter shaded area).

## References

1. Bliss, T.V., and Collingridge, G.L. (1993). A synaptic model of memory: long-term potentiation in the hippocampus. Nature 361, 31–39.

2. Lau, C.G., and Zukin, R.S. (2007). NMDA receptor trafficking in synaptic plasticity and neuropsychiatric disorders. Nat Rev Neurosci 8, 413–426.

3. Paoletti, P., Bellone, C., and Zhou, Q. (2013). NMDA receptor subunit diversity: impact on receptor properties, synaptic plasticity and disease. Nat Rev Neurosci 14, 383–400.

4. Zukin, R.S., and Bennett, M.V. (1995). Alternatively spliced isoforms of the NMDARI receptor subunit. Trends Neurosci 18, 306–313.

5. Nakanishi, N., Axel, R., and Shneider, N.A. (1992). Alternative splicing generates functionally distinct N-methyl-D-aspartate receptors. Proc Natl Acad Sci U S A 89, 8552–8556.

6. Laurie, D.J., and Seeburg, P.H. (1994). Regional and developmental heterogeneity in splicing of the rat brain NMDAR1 mRNA. J Neurosci 14, 3180–3194.

7. Paupard, M.C., Friedman, L.K., and Zukin, R.S. (1997). Developmental regulation and cell-specific expression of N-methyl-D-aspartate receptor splice variants in rat hippocampus. Neuroscience 79, 399–409.

8. Kuppenbender, K.D., Albers, D.S., Iadarola, M.J., Landwehrmeyer, G.B., and Standaert, D.G. (1999). Localization of alternatively spliced NMDAR1 glutamate receptor isoforms in rat striatal neurons. J Comp Neurol 415, 204–217.

9. Nash, N.R., Heilman, C.J., Rees, H.D., and Levey, A.I. (1997). Cloning and localization of exon 5-containing isoforms of the NMDAR1 subunit in human and rat brains. J Neurochem 69, 485–493.

10. Rumbaugh, G., Prybylowski, K., Wang, J.F., and Vicini, S. (2000). Exon 5 and spermine regulate deactivation of NMDA receptor subtypes. J Neurophysiol 83, 1300–1306.

11. Vance, K.M., Hansen, K.B., and Traynelis, S.F. (2012). GluN1 splice variant control of GluN1/GluN2D NMDA receptors. J Physiol 590, 3857–3875.

12. Liu, H., Wang, H., Peterson, M., Zhang, W., Hou, G., and Zhang, Z.W. (2019). N-terminal alternative splicing of GluN1 regulates the maturation of excitatory synapses and seizure susceptibility. Proc Natl Acad Sci U S A 116, 21207–21212.

13. Traynelis, S.F., Hartley, M., and Heinemann, S.F. (1995). Control of proton sensitivity of the NMDA receptor by RNA splicing and polyamines. Science 268, 873–876.

14. Sengar, A.S., Li, H., Zhang, W., Leung, C., Ramani, A.K., Saw, N.M., Wang, Y., Tu, Y., Ross, P.J., Scherer, S.W., et al. (2019). Control of Long-Term Synaptic Potentiation and Learning by Alternative Splicing of the NMDA Receptor Subunit GluN1. Cell Reports 29, 4285–4294.e4285.

15. Durand, G.M., Gregor, P., Zheng, X., Bennett, M.V., Uhl, G.R., and Zukin, R.S. (1992). Cloning of an apparent splice variant of the rat N-methyl-D-aspartate receptor NMDAR1 with altered sensitivity to polyamines and activators of protein kinase C. Proc Natl Acad Sci U S A 89, 9359–9363.

16. Horak, M., and Wenthold, R.J. (2009). Different roles of C-terminal cassettes in the trafficking of full-length NR1 subunits to the cell surface. J Biol Chem 284, 9683–9691.

17. Lee, J.A., Xing, Y., Nguyen, D., Xie, J., Lee, C.J., and Black, D.L. (2007). Depolarization and CaM kinase IV modulate NMDA receptor splicing through two essential RNA elements. PLoS Biol 5, e40.

18. Mu, Y., Otsuka, T., Horton, A.C., Scott, D.B., and Ehlers, M.D. (2003). Activity-dependent mRNA splicing controls ER export and synaptic delivery of NMDA receptors. Neuron 40, 581–594.

19. Okabe, S., Miwa, A., and Okado, H. (1999). Alternative splicing of the C-terminal domain regulates cell surface expression of the NMDA receptor NR1 subunit. J Neurosci 19, 7781–7792.

20. Scott, D.B., Blanpied, T.A., Swanson, G.T., Zhang, C., and Ehlers, M.D. (2001). An NMDA receptor ER retention signal regulated by phosphorylation and alternative splicing. J Neurosci 21, 3063–3072.

21. Hansen, K.B., Yi, F., Perszyk, R.E., Furukawa, H., Wollmuth, L.P., Gibb, A.J., and Traynelis, S.F. (2018). Structure, function, and allosteric modulation of NMDA receptors. J Gen Physiol 150, 1081–1105.

22. Traynelis, S.F., Wollmuth, L.P., McBain, C.J., Menniti, F.S., Vance, K.M., Ogden, K.K., Hansen, K.B., Yuan, H., Myers, S.J., and Dingledine, R. (2010). Glutamate receptor ion channels: structure, regulation, and function. Pharmacol Rev 62, 405–496.

23. Rajani, V., Sengar, A.S., and Salter, M.W. (2020). Tripartite signalling by NMDA receptors. Molecular Brain 13.

24. Nong, Y., Huang, Y.-Q., Ju, W., Kalia, L.V., Ahmadian, G., Wang, Y.T., and Salter, M.W. (2003). Glycine binding primes NMDA receptor internalization. Nature 422, 302–307.

25. Han, L., Campanucci, V.A., Cooke, J., and Salter, M.W. (2013). Identification of a single amino acid in GluN1 that is critical for glycine-primed internalization of NMDA receptors. Molecular Brain 6, 36.

26. Parikshak, N.N., Swarup, V., Belgard, T.G., Irimia, M., Ramaswami, G., Gandal, M.J., Hartl, C., Leppa, V., Ubieta, L.D.L.T., Huang, J., et al. (2016). Genome-wide changes in lncRNA, splicing, and regional gene expression patterns in autism. Nature 540, 423–427.

27. Adie, E.J., Kalinka, S., Smith, L., Francis, M.J., Marenghi, A., Cooper, M.E., Briggs, M., Michael, N.P., Milligan, G., and Game, S. (2002). A pH-sensitive fluor, CypHer 5, used to monitor agonist-induced G protein-coupled receptor internalization in live cells. Biotechniques 33, 1152–1154, 1156–1157.

28. Nong, Y., Huang, Y.Q., and Salter, M.W. (2004). NMDA receptors are movin’ in. Curr Opin Neurobiol 14, 353–361.

29. Macia, E., Ehrlich, M., Massol, R., Boucrot, E., Brunner, C., and Kirchhausen, T. (2006). Dynasore, a cell-permeable inhibitor of dynamin. Dev Cell 10, 839–850.

30. Huntley, M.A., Srinivasan, K., Friedman, B.A., Wang, T.M., Yee, A.X., Wang, Y., Kaminker, J.S., Sheng, M., Hansen, D.V., and Hanson, J.E. (2020). Genome-Wide Analysis of Differential Gene Expression and Splicing in Excitatory Neurons and Interneuron Subtypes. J Neurosci 40, 958–973.

31. Raeder, H., Holter, S.M., Hartmann, A.M., Spanagel, R., Moller, H.J., and Rujescu, D. (2008). Expression of N-methyl-d-aspartate (NMDA) receptor subunits and splice variants in an animal model of long-term voluntary alcohol self-administration. Drug Alcohol Depend 96, 16–21.

32. Regan, M.C., Grant, T., McDaniel, M.J., Karakas, E., Zhang, J., Traynelis, S.F., Grigorieff, N., and Furukawa, H. (2018). Structural Mechanism of Functional Modulation by Gene Splicing in NMDA Receptors. Neuron 98, 521–529 e523.

33. Saito, Y., Yuan, Y., Zucker-Scharff, I., Fak, J.J., Jereb, S., Tajima, Y., Licatalosi, D.D., and Darnell, R.B. (2019). Differential NOVA2-Mediated Splicing in Excitatory and Inhibitory Neurons Regulates Cortical Development and Cerebellar Function. Neuron 101, 707–720 e705.

34. Ferreira, J.S., Papouin, T., Ladépêche, L., Yao, A., Langlais, V.C., Bouchet, D., Dulong, J., Mothet, J.-P., Sacchi, S., Pollegioni, L., et al. (2017). Co-agonists differentially tune GluN2B-NMDA receptor trafficking at hippocampal synapses. eLife 6.

35. Li, L.-J., Hu, R., Lujan, B., Chen, J., Zhang, J.-J., Nakano, Y., Cui, T.-Y., Liao, M.-X., Chen, J.-C., Man, H.-Y., et al. (2016). Glycine Potentiates AMPA Receptor Function through Metabotropic Activation of GluN2A-Containing NMDA Receptors. Frontiers in Molecular Neuroscience 9.

